# Loss of multiple micro-RNAs uncovers multi-level restructuring of gene regulation in rodents

**DOI:** 10.1101/2024.12.05.627021

**Authors:** Felix Langschied, Matthias S. Leisegang, Stefan Günther, Fabian Hahner, Ralf P. Brandes, Ingo Ebersberger

## Abstract

**Background:** The regulatory network that coordinates gene expression ultimately determines the phenotype of an organism. Micro-RNAs (miRNAs) are post-transcriptional regulators involved in key biological processes. Lineage-specific losses of multiple miRNA families are rare, and reported cases of multiple miRNA losses coincide with significant changes in gene regulation resulting in body plan modifications. Recently, 15 mammalian miRNA families were found to be missing in the *Eumuroidea*, the rodent lineage that includes the model organisms mouse and rat. However, the impact of their absence on the gene regulatory networks in this lineage remains unknown.

**Results:** The *in silico* characterization of all 15 miRNAs revealed that their absence is best explained by multiple independent losses. Analyzing their target genes in humans reveals a significant enrichment of GO-terms linked to cellular and developmental processes. Overexpressing two of the co-absent miRNAs, Mir-197 and Mir-769, in human and mouse inducible pluripotent stem cells (iPSCs) resulted in significantly perturbed expression patterns in both species. *In silico* target site prediction revealed a significant enrichment of direct targets exclusively in the down-regulated genes. Four genes were down-regulated in both mouse and human and maximum parsimony suggests that the corresponding miRNA target sites were already present in the last common ancestor of mammals. The response of these genes to miRNA overexpression in mice, therefore, unveils remnants of the ancient gene regulatory network that have persisted until today. The evolutionary age of these regulatory connections provides initial evidence that the miRNA losses in *Eumuroidea* must have had consequences for the regulation of gene expression. Notably, we show that the miRNA loss in the Eumuroidea coincides with the largest lineage-specific loss of transcription factors within the mammals.

**Conclusions:** The impact of miRNA losses has thus far been investigated on a gene-by-gene basis. Our findings indicate that cooperative effects between miRNAs should be considered when assessing the impact of miRNA loss. We provide evidence that the *Eumuroidea* have modified their gene regulatory networks on two levels, transcriptionally and post-transcriptionally. It will now be interesting to precisely chart the differences in gene regulation and assess their combined impact on the suitability of mice and rats as model systems for human disease.

## Background

Micro-RNAs (miRNAs) are post-transcriptional regulators, that each downregulate multiple mRNAs (1). The targeting process is mainly guided by the pairing of a 6-8 nt long seed sequence in the miRNA with a matching target site in the mRNA (2). Due to the short length of the seed sequences, matching target sites are likely to occur in many mRNAs by chance. It has therefore been proposed that young miRNAs initially target a random set of genes (3). If the knockdown of any initial targets is deleterious, two possible outcomes may occur. Either the new miRNA is removed from the population (4), or the deleterious target sites degrade, allowing the novel miRNA gene to persist (5). Once a miRNA has become fixed in a population, its target specificity increases over time (6). Additionally, miRNAs tend to acquire more targets over time (7), coinciding with their expression in an increasing number of tissues (4). This aligns well with the effect size of miRNA knockouts. Evolutionary old miRNAs that emerged prior to vertebrate diversification assume key roles in cellular regulatory networks, and the knockout of individual miRNA genes often results in lethal phenotypes (8). In contrast, the knock-out of an evolutionarily younger miRNA, which can be traced back to the last common ancestor (LCA) of placental mammals, generally has only a modest effects, such as growth defects (9) or anxiety-related behaviour (10). However, what remains largely unknown is the extent to which regulatory networks are shaped by a cooperative effect of multiple younger miRNAs.

In the context of proteins, cooperative effects can be identified if the corresponding genes share a similar presence-absence pattern across taxon collections (11). These patterns, also known as phylogenetic profiles, help in identifying genes that jointly establish, e.g., a metabolic pathway (12), or a protein complex (13). In such cases, the loss of one gene can abolish the function of the entire pathway/complex, if it is not compensated for. This makes it likely that the remaining proteins become dispensable, and their subsequent loss leaves a characteristic footprint in their phylogenetic profiles. In turn, the analysis of even vague annotations of functionally integrated genes can provide a mosaic-like view of the overarching function they convey (14). To date, it remains unknown whether concerted loss is also a signal of functional integration for miRNAs, or even how an overlap in miRNA function might manifest itself.

Over evolutionary time scales, miRNA families shared across multiple, taxonomically diverse species are generally resistant to gene loss (15,16). However, there are a few exceptions, most notably in lineages that have significantly simplified their body plans, such as flatworms and sea squirts (17,18). In mammals, an analysis of eight species identified the loss of several miRNAs in mouse and rat that cannot be attributed to body plan simplification (18). However, the limited taxonomic resolution of that study makes it difficult to determine when these losses occurred during the rodent diversification.

Consequently, connecting the miRNA loss to phenotypic changes has remained challenging, obscuring the functional consequences of the corresponding alterations in the gene regulatory network.

To shed further light on how the concerted loss of multiple miRNAs impacts the regulatory network, we focused on 15 mammalian miRNA families that we previously reported as lost during rodent diversification (16). We hypothesized that these losses signal a restructuring of the gene regulatory network. To investigate this, we first characterized the evolution and likely function of the absent miRNA families *in silico*. Overexpression of two miRNAs in human and murine inducible pluripotent stem cells (iPSCs) provided insights into primordial regulatory connections in the LCA of human and mouse that were likely disrupted by the miRNA loss on the rodent lineage. Scanning the entire protein-coding gene set of humans revealed an enrichment of transcription factors among the genes co-lost with the miRNA families in the *Eumuroidea*.

## Methods

### Phylogenetic profiles

miRNA phylogenetic profiles were constructed as described previously (16). Briefly, we run ncOrtho in default settings to train covariance models (19) for all human miRNA genes annotated in MirGeneDB 2.0 (20), collecting training data in four primate genomes (*Macaca mulatta, Gorilla gorilla*, *Pongo abelii* and *Nomascus leucogenys).* The covariance models are then used in the second step of ncOrtho to search for miRNA orthologs in 169 Eutherian genomes annotated in RefSeq NCBI Refseq Genomes release 207 (21). The targeted ortholog search tool fDOG (22) was used to construct phylogenetic profiles of all human protein-coding genes across all taxa for which miRNA ortholog assignments by ncOrtho are available. Phylogenetic profiles of all protein-coding genes and miRNA genes from the 16 families lost in Eumuroidea were represented as binary vectors. miRNA orthologs with non-conserved seed sequence were treated as ‘absent’ since they likely have changed in function. The top percentile of protein-coding genes with the highest Pearson correlation coefficient to each miRNA profile were considered to have similar phylogenetic profiles. Then, genes whose profile was similar to at least half of the lost miRNAs were identified as sharing an evolutionary history with the lost miRNA families.

### miRNA targets

Experimentally validated targets of all 15 lost miRNAs were downloaded from MirTarBase 9 (23) and filtered for interactions with strong experimental evidence. Additionally, targets of Mir-197 and Mir- 769 were predicted with TargetScan7 (2). TargetScan provides the option to predict target sites using the context++-score method which is independent of the conservation status of the miRNA in 10 vertebrate species. We calculated the total unweighted context++-score (tc++) as described in Agarwal et al. (2015) and set a minimum tc++ threshold of -0.2 to define predicted targets (Supplementary Figure S1). Overlapping target sites in the 3’-UTRs of orthologous genes were considered homologous, and we used this information to date the emergence of these sites. The most parsimonious branch of target site origin was determined using the Count software package (24).

### Cell culture

Human induced pluripotent stem cells (iPSCs) (WSTIi081-A, EbiSC) were cultured in TeSR™-E8™ medium (#05990, STEMCELL™ Technologies) at 37°C and 5% CO2 in a humidified atmosphere. Mouse embryonic fibroblasts (MEF) were isolated from C57B6 mice and cultured as described previously (25). Isolated mouse embryonic fibroblasts were reprogrammed into induced pluripotent stem cells (iPSCs) using the STEMCCA system (26). Briefly, 20.000 mouse embryonic fibroblast (were seeded into culture dishes. After 24 h, cells were infected with STEMCCA virus for 24 h. Media was changed for five consecutive days. Cells were then reseeded on mitomycin (Serva, 10 μg/mL, 3 h) -inactivated feeder cells until murine induced pluripotent colonies emerged. Individual colonies were subsequently picked and expanded. Cells were passaged (DMEM GlutaMAX, #10566016, Gibco; 20 % Knockout Serum Replacement, #10828028, 25 μg mLIF, #250–02, Peprotech; 0.1% β-mercaptoethanol, 200 μM Glutamine, #11539876, Gibco) by 70% before colonies touched and reseeded between 2.5 × 10^3^ and 5 × 10^3^ cells per cm^2^.

### FACS sorting of mouse iPSCs

Mouse induced pluripotent stem cells (iPSC) cultured on feeder cells (MEF) or only feeder cells (negative control) were dissociated with accutase (#A6964, Sigma-Aldrich) centrifuged and resuspended to single cell suspension in FACS Buffer (PBS, 5 % FCS). Cells were sorted using the Sony SH800 at 4°C (nozzle size 100 um, flow rate 6.0). Cells were selected using forward scatter and back scatter (4, 25 %). Gating was done by using autofluorescence of cells (FITC-A-Compensated vs. PE-A-Compensated). Feeder used as negative control to differentiate between cell types in mixed culture of feeder and miPSC. Gating was set to sort for miPSC (100.000 events).

### Transfection of human and murine iPSCs with miRNA precursors

For miRNA overexpression treatments, 250.000 human or murine iPSCs were transfected with Lipofectamin RNAiMAX (Cat. No. 13778075) according to the manufacturers protocol (ThermoFisher Scientific). Ambion® Pre-miR™ miRNA Precursor for hsa-miR-197-3p (Assay ID PM10354, Cat. No. AM17100, ThermoFisher Scientific) and hsa-miR-769-5p (Assay ID PM11974, Cat. No. AM17100, ThermoFisher Scientific) were used. As negative control, Pre-miR™ miRNA Precursor Negative Control #1 (Cat. No. AM17110, ThermoFisher Scientific) was used. All miRNA precursor transfections were performed with a concentration of 8 nM for 48 h.

### RNA isolation, Reverse transcription and RT-qPCR

Total RNA isolation was performed with the RNA Mini Kit (Bio&Sell) according to the manufacturers protocol and reverse transcription was performed with the SuperScript III Reverse Transcriptase (Thermo Fisher) using a combination of oligo(dT)23, random hexamer as well as U6 snRNA- and miRNA-specific RT primers (Sigma). The resulting cDNA was amplified in an AriaMX cycler (Agilent) with the ITaq Universal SYBR Green Supermix and ROX as reference dye (Bio-Rad, Cat. No. 1725125). Relative expression of human target genes was normalized to U6 snRNA. Expression levels were analyzed by the delta-delta Ct method with the AriaMX qPCR software (Agilent, Version 1.7). The primers for the detection of miRNAs were designed with miRprimer2 (27). The following oligonucleotide sequences were used for the miRNAs: miR-197-5p RT 5’-GTC GTA TCC AGT GCA GGG TCC GAG GTA TTC GCA CTG GAT ACG ACC CTC CC-3’, forward 5’- GCG GCG GCG GGT AGA GAG GGC AG-3’; miR-197-3p RT 5’-GTC GTA TCC AGT GCA GGG TCC GAG GTA TTC GCA CTG GAT ACG ACG CTG GG-3’, forward 5’-GCG GCG GTT CAC CAC CTT CTC C-3’; miR-769-5p RT 5’-GTC GTA TCC AGT GCA GGG TCC GAG GTA TTC GCA CTG GAT ACG ACA GCT CA-3’, forward 5’-GCG GCG GTG AGA CCT CTG GGT TC-3’; miR-769-3p RT 5’-GTC GTA TCC AGT GCA GGG TCC GAG GTA TTC GCA CTG GAT ACG ACA ACC AA-3’, forward 5’-GCG GCG GCT GGG ATC TCC GGG G-3’. For all miRNAs, a universal reverse primer was used: 5’-ATC CAG TGC AGG GTC CGA GG-3’. For U6 snRNA, the following oligonucleotides were used: RT 5’-CGC GCC TGC AGG TCG ACA ATT AAC CCT CAC TAA AGG GTT GCG TGT CAT CC-3’, forward 5’-GTA ATA CGA CTC ACT ATA GGG AGA AGA GCC TGC GCA AGG-3’ and reverse 5’-CAG GTC GAC AAT TAA CCC TC-3’.

### RNA-Seq

RNA amounts were normalized and 500-100ng of total RNA was used as input for SMARTer Stranded Total RNA Sample Prep Kit HI Mammalian (Takara Bio). Sequencing was performed on the NextSeq2000 platform (Illumina) using P3 flowcell with 72bp single-end setup. Trimmomatic version 0.39 was employed to trim reads after a quality drop below a mean of Q15 in a window of 5 nucleotides and keeping only filtered reads longer than 15 nucleotides (28). Reads were aligned versus Ensembl human genome version hg38 (release 104) or Ensembl mouse genome version mm39 (Ensembl release 104) with STAR 2.7.9a (29). Alignments were filtered with Picard 2.25.2 (30) to remove duplicates, as well as multi-mapping, ribosomal, or mitochondrial reads. Gene counts were established with featureCounts 2.0.2 by aggregating reads overlapping exons on the correct strand excluding those overlapping multiple genes (31). The raw count matrix was normalized with DESeq2 version 1.30.0 (32).

## Results

### Characterization of miRNA loss and test for functional compensation

miRNAs are rarely lost in vertebrates, but there are a few lineage-specific exceptions, with the absence of 15 miRNA families in rodents being among the most prominent one (16). We first ensured that the observed losses were not inferred due to sensitivity limits of the ortholog search or assembly artefacts. Consulting the 100-vertebrate whole genome alignment in the UCSC Genome Browser (33) revealed mutational changes ranging from single nucleotide substitutions to small scale insertion/deletion events in the miRNA loci (Figure 1B, Supplementary Figure S2). To determine whether few or even a single large-scale event could explain the observed losses, we analysed the position of the miRNA genes in the human genome. The corresponding genes are spread across multiple loci (**Error! Reference source not found.**), strongly suggesting that multiple independent losses account for the absence of these 15 miRNA families. To determine when the loss events leading to the shared absence of these families occurred, we performed an in-depth characterization of their phylogenetic profiles. This indicated that the respective miRNA genes were lost at different points during the diversification of the *Glires* (*Rodentia* and *Lagomorpha*) (Figure 1A). Notably, some of these miRNA families were also independently lost in the insectivores (*Eulipotyphla*) and inside the bats (*Chiroptera*; Figure 1A).

**Figure 1:**
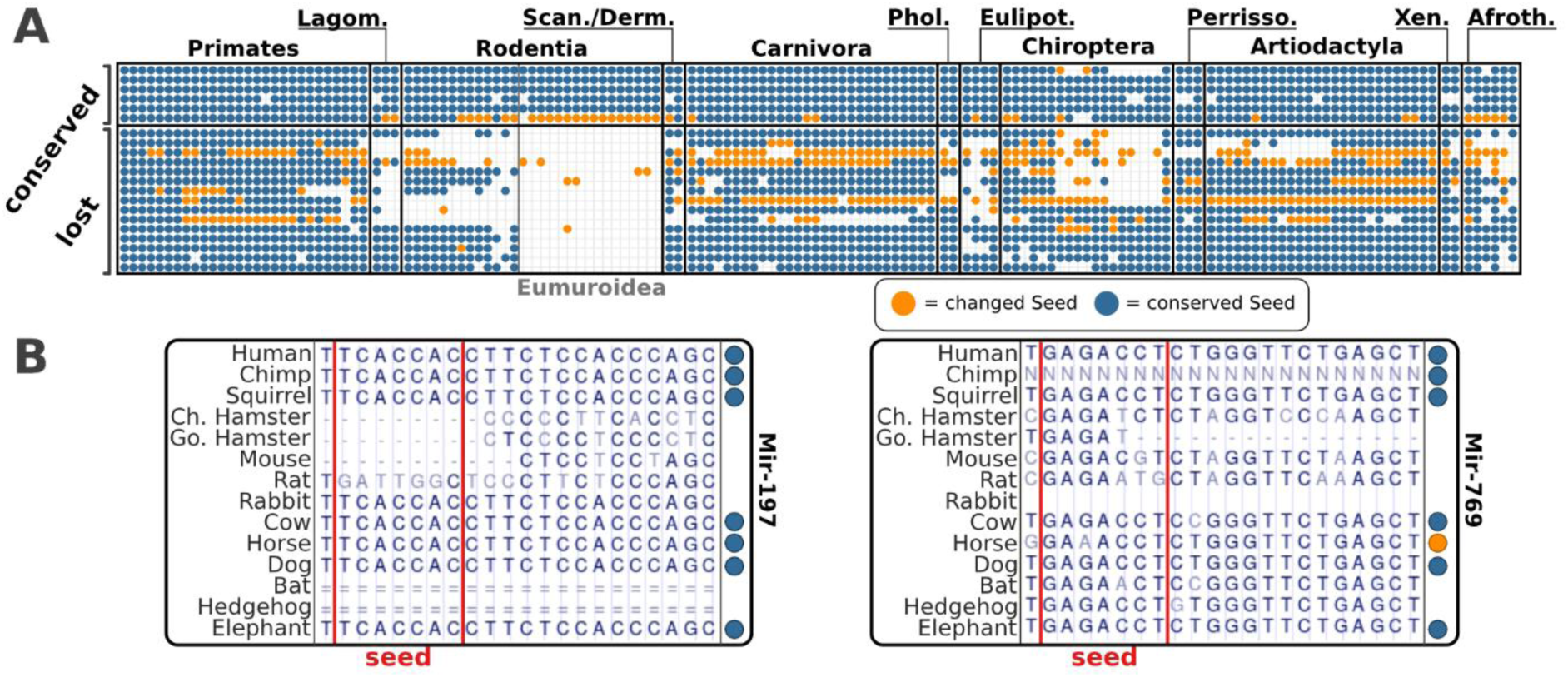
Characterization of miRNA families co-absent in the *Eumuroidea*. A) Phylogenetic profiles of 21 miRNA families of which the top six are conserved throughout the vertebrates and the remaining 15 have been lost in the Rodentia. A dot indicates that at least one ortholog of any pre-miRNA of the miRNA family was identified in the respective species. Dot color shows whether the human seed sequence was found in the ortholog (blue) or not (orange). Species are ordered according to increasing taxonomic distance to the reference species of the ortholog search (human) from left to right. Lagom. = Lagomorpha, Scan. = Scandentia, Derm. = Dermoptera, Phol. = Pholidota, Eulipot. = Eulipothyphla, Perrisso. = Perrissodactyla, Xen. = Xenarthra, Afroth. = Afrotheria. B) 14-species alignment for the genomic loci harboring miRNA families 197 and 769 in humans extracted from the 100-vertebrate whole genome alignments. The alignments confirm the loss of the miRNA in *Eumuroidea*.

We next investigated whether the absence of the miRNA families is compensated by either redundant or newly emerged miRNAs. Since seed-pairing is the primary determinant of miRNA targeting specificity (2), we first confirmed that the seed sequences of the co-absent miRNA families are not shared by any of the rodent-specific miRNAs annotated in MirGeneDB (34). To increase the sensitivity of the analysis, we subsequently used each of the 15 mature miRNA sequences as a query for a BLASTn-short search in the collection of mature mouse and rat miRNAs in MirGeneDB. This obtained no significant hit (E-value < 0.01) indicating that there are no rodent-specific miRNAs that compensate for the observed loss using the same target sites.

### Target genes of co-absent miRNAs are involved in the regulation of developmental processes

To identify potential functional consequences of the reduced miRNA set in *Eumuroidea*, we analyzed publicly available datasets of miRNA tissue expression (34) and experimentally validated target genes (23). Given the collective absence of 15 miRNA families in *Eumuroidea*, it is tempting to speculate about the functional integration of the respective miRNA genes. To investigate this, we first analyzed whether the co-absent miRNA genes exhibit tissue-specific expression. A comparison of their expression levels across 45 human tissues revealed that the expression vectors of the co-absent miRNA genes are not more similar than expected by chance (Supplementary Figure S3). Hence, gene expression across tissues provides no complementary evidence for the functional integration of these genes. We next collected experimentally validated target genes recorded in the miRTarBase database (23). Out of the 85 recorded regulatory connections, 50 belong to the Mir-506 family, while the remaining 35 are distributed among seven other miRNA families (**Error! Reference source not found.**). We found no significant overlap between these sets of target genes, indicating no evidence for functional integration of co-absent miRNAs via shared target catalogues (Supplementary Figure S4). However, the combined set of 85 target genes was significantly enriched with Gene Ontology (GO) terms associated with the regulation of cellular and developmental processes (Supplementary Figure S5). This provides an initial indication that there is a functional overlap of genes targeted by the co-absent miRNAs.

### miRNA overexpression affects similar numbers of genes irrespective of whether the host organism contains a native copy

All 15 miRNA families date back at least to the LCA of placental mammals (see Fig. 1). It is therefore conceivable that their regulatory networks were already established when the mouse and human lineages split. Expressing human miRNA orthologs in mouse should allow investigating if and to what extent this regulatory network has altered after the loss, and whether its remnants can still be traced in rodents. We selected Mir-197 and Mir-769 as candidates for the experiments as these were single-copy miRNA families with moderate expression levels under native conditions in human (Supplementary Figure S6). An overexpression of the corresponding miRNAs should therefore impose a noticeable effect on the target genes’ expression even in humans that contain a native copy. To test this, we performed differential gene expression analyses after overexpression of the two human miRNAs in both human and mouse iPSCs. While truly comparable cell lines of organ origin do not exist between human and mice (35), naïve-state iPSCs should be more similar between species than differentiated cell types. In response to overexpression, the abundance of the mature miRNAs Mir- 197-3p and Mir-769-5p significantly increased in both human and murine iPSCs (Figure 2A, B). To reveal the effect of this overexpression on a transcriptome-wide level, RNA-Seq was performed in both human and murine iPSCs. The replicates from the four treatment conditions and the two controls form clusters in a principal component analysis of the normalized expression values, indicating a condition-specific impact of miRNA overexpression in both organisms (Figure 2C).

**Figure 2:**
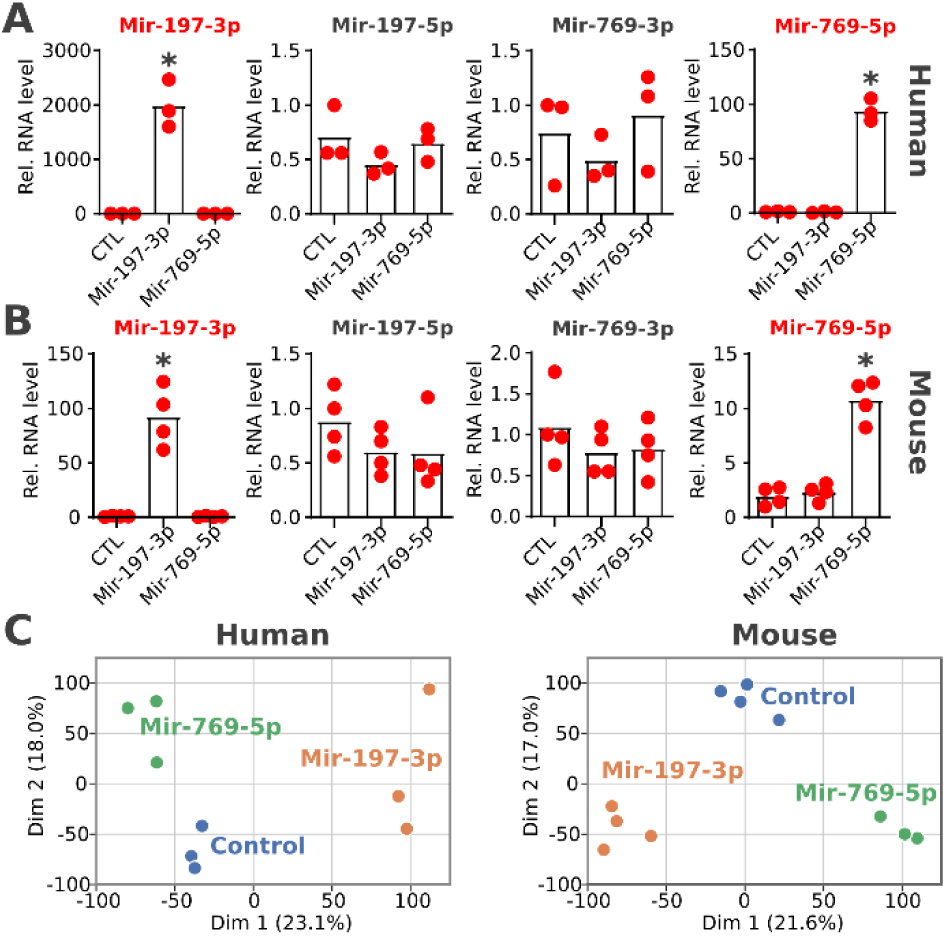
Effect of overexpression of human pre-miRNAs in human and murine iPSCs. RNA levels relative to U6 snRNA are shown for both miRNAs and the miRNA Precursor Negative Control (CTL) in both human (A, n=3) and murine iPSCs (B, n=4). The name of the mature miRNA is highlighted in red; the name of the star miRNA is shown in black. Significantly altered expression levels are indicated with an asterisk (unpaired t-test, p-value < 0.01). C) Principal component analysis of normalized expression values after RNA-seq.

In human iPSCs, 194 genes (Mir-197) and 103 genes (Mir-769) showed significant changes in their expression levels in response to miRNA overexpression (Table 1, **Error! Reference source not found.**). In mice, the heterologous expression of the human miRNAs resulted in a significant change of expression levels for 216 (Mir-197) and 228 (Mir-769) genes. This confirms the expectation that the overexpression of the two miRNAs has a substantial effect on gene expression in human iPSCs. More importantly, this shows that the mouse transcriptome responds to the presence of the human miRNAs in the same order of magnitude.

**Table 1:** Differentially expressed genes after miRNA overexpression. Protein-coding genes were considered differentially expressed if their mRNAs have an absolute log2fold-change > 0.5, at least 100 reads in the negative control and an adjusted p-value <= 0.05 (FDR 5%)

| Organism | miRNA | Up | Down |
| --- | --- | --- | --- |
| Human | Mir-197-3p | 85 | 109 |
|  | Mir-769-5p | 25 | 78 |
| Mouse | Mir-197-3p | 101 | 115 |
|  | Mir-769-5p | 76 | 152 |

As expected from the mode of miRNA activity, most genes that responded to miRNA overexpression were downregulated in both cell lines and in both treatments. However, a considerable number of genes also increased their expression levels, which cannot be explained as a direct effect of the miRNA overexpression. Instead, upregulated genes indicate the presence of indirect effects on the regulatory network. Indeed, we found between 6 and 8 down-regulated transcription factors (TFs) in each treatment condition, which could mediate this effect (**Error! Reference source not found.**; Lambert et a l. 2018). It is conceivable that the miRNA-driven downregulation of these TFs propagates regulatory effects to further genes that are not directly targeted by the miRNA. Since these indirect effects can also induce down-regulation of genes, mimicking the mode of action of the miRNA, we next identified the most likely primary targets of the two miRNAs in human and murine iPSCs.

### Target prediction identifies physiologically relevant targets of lost miRNAs in both human and mouse

To identify primary targets of Mir-197 and Mir-769 in human and murine iPSCs, *in silico* target prediction was performed using the context++-score method implemented in TargetScan 7 (2). This predicted 1079 genes as potential targets of Mir-197 in human, and 792 targets in mouse. For Mir-769, TargetScan 7 predicted 935 and 839 targets for human and mouse, respectively. Notably, the genes we found up-regulated in the presence of the miRNAs contain no more targets than expected by chance. This is in line with the hypothesis that their expression change is an indirect effect (Figure 3B). In contrast, down-regulated genes were significantly enriched with predicted targets, which aligns with the expectation given the mode of action of miRNAs. The four sets of genes that are down-regulated as well as predicted as targets are thus considered primary targets of Mir-197 and Mir-769 in human or murine iPSCs, respectively. Notably, none of the experimentally validated targets listed in miRTarBase overlapped with the primary targets of the two miRNAs, as none of the miRTarBase targets are expressed in human iPSCs (**Error! Reference source not found.**, **Error! Reference source not found.**).

**Figure 3:**
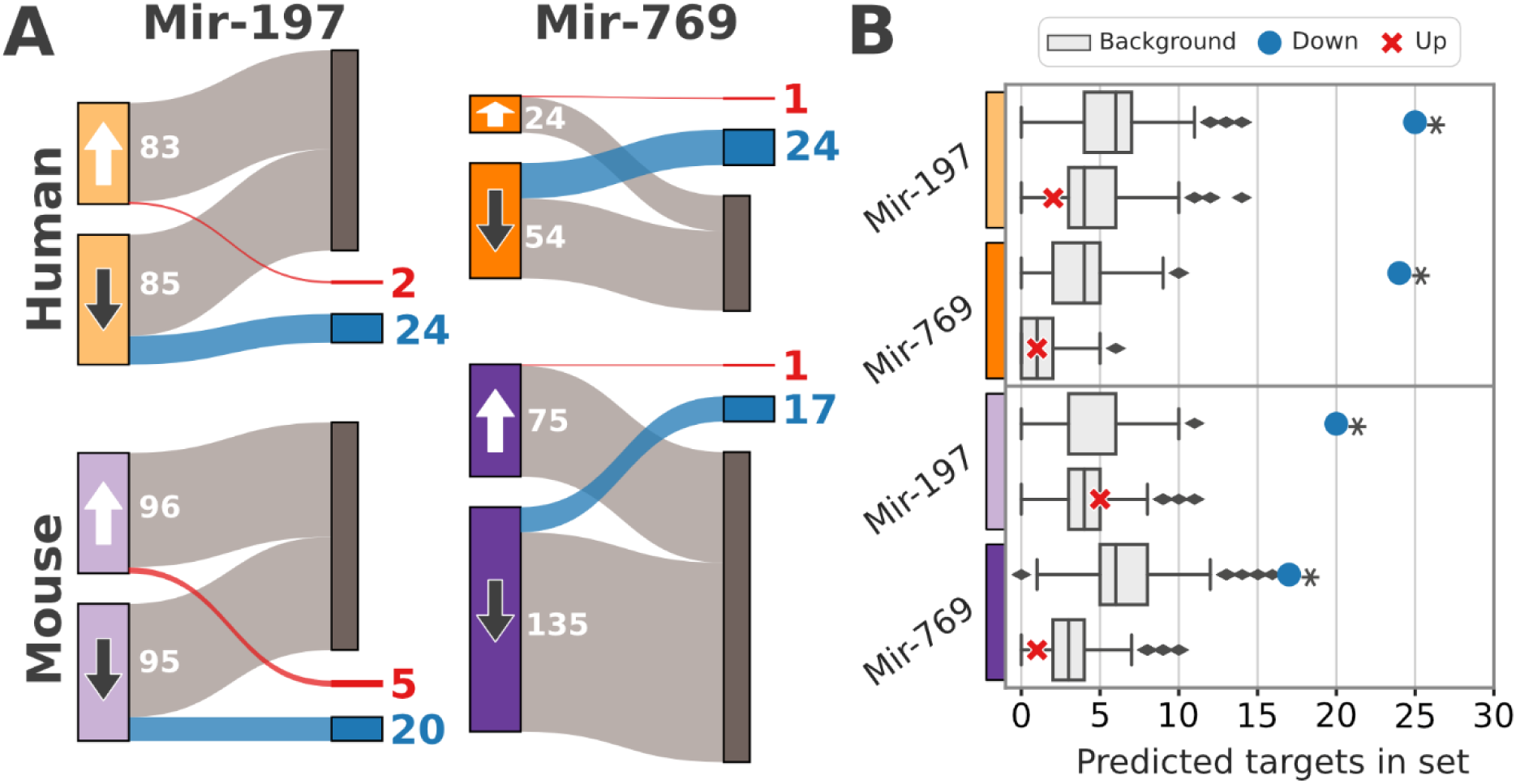
Overlap between predicted miRNA targets and differentially expressed genes. A) Sets of significantly up- and down-regulated genes are marked with white and dark arrows, respectively. Overlaps with predicted targets are marked in red for up-regulated genes and blue for down-regulated genes. B) Background distribution for the number of predicted targets was created by sampling 1,000 gene sets matching the number of up- or down-regulated transcripts for each miRNA and species. Then, genes whose mRNAs are predicted as targets were counted and compared to the observed number. Observations that deviate significantly from the background distribution are marked with an asterisk (two-sided Z-score test; p-value <= 0.01).

### Overlap of primary miRNA targets in human and murine cells indicate remnants of ancient regulatory connections

Our analysis has identified four sets of 17 to 24 primary targets in human or murine iPSCs for each miRNA. No gene was shared between the sets for Mir-197 and Mir-769, which is in line with our previous analysis based on targets provided by miRTarBase. Consulting GO-annotations as well as available information about protein interactions (37) provided no statistically significant functional overlap between targets of the two miRNAs. We next pursued the orthogonal approach and compared the set of primary targets for the same miRNA between human and mouse. There are four genes, ARMC1, ATP6V1A, CCDC85C and TTPAL that are primary targets of Mir-197 in both human and murine iPSCs (Figure 4A). For Mir-769, no shared target genes met the selected filter criteria, however several genes just fell below the inclusion threshold (**Error! Reference source not found.**). If we slightly lower t he minimum log2fold-change to 0.25, we identified an overlap of three genes between human and murine iPSCs for Mir-769 (**Error! Reference source not found.**).

**Figure 4:**
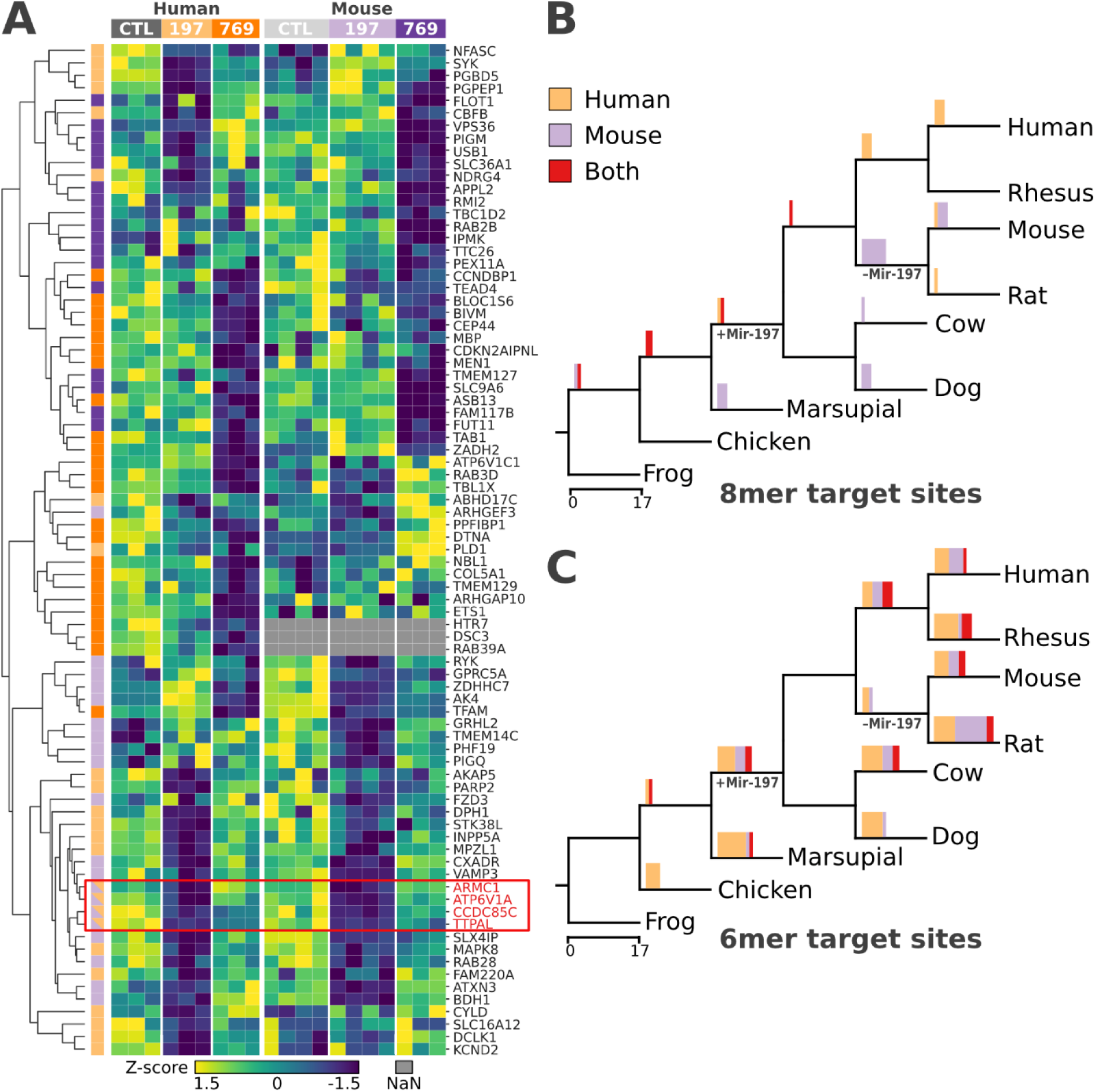
Primary targets in human and murine iPSCs partly overlap due to evolutionary old target sites. **A)** Z-score of normalized read counts between the negative control (CTL) and the overexpression of Mir-197-3p or Mir-769-5p per organism. Rows are clustered by cosine similarity. The condition in which a gene is downregulated as well as predicted as a target for the respective miRNA are indicated with colored boxes on the left. The color code follows the top boxes where targets of Mir-197 and Mir- 769, in human are colored in light orange and orange, respectively. Murine targets are colored in light purple for Mir-197 and purple for Mir-769. The red box highlights the four genes that are significantly down-regulated and predicted as targets for Mir-197 in both human and mouse. Gene labels follow the human nomenclature, all names of the mouse orthologs can be derived by capitalization. B) Dating the emergence of target sites in species-specific and overlapping targets of Mir-197. Bar charts describe the number of target sites gained in the node to the right of the edge. The color of the bars indicates whether the corresponding target gene is downregulated in human (orange), mouse (purple), or both (red). C) Origin of 6mer target sites. Color code follows B).

An overlap of primary targets of Mir-197 in human and mouse could be either a remnant of the ancient regulatory network, or the result of younger target sites gained independently in the human or rodent lineage. To differentiate between these scenarios, we dated the emergence of target sites in all primary targets of Mir-197 (Figure 4B, C). This revealed that 8mer target sites of genes that respond to the overexpression of Mir-197 in only one of the two species emerged after the divergence of the human and mouse lineages (Figure 4B). Consequently, the regulatory network of Mir-197 appears to be highly flexible. In contrast, 8mer target sites in genes that are primary targets in both human and mouse are at least as old as the LCA of the two species (Figure 4B). We find no such signal when repeating the analysis with 6mers (Figure 4C). This is consistent with 6mer target sites having a markedly less pronounced impact on transcript abundance than 8mer sites (1). They are consequently more likely to be gained (and lost) as a stochastic effect of sequence evolution than 8mer target sites.

Taken together, our findings suggest that at least Mir-197 was part of an ancient regulatory network in the LCA of human and mouse. Individual ancient regulatory connections have been conserved on both evolutionary lineages such that Mir-197 still conveys a physiologically measurable effect on transcript abundance in contemporary human and mouse. This strongly indicates that Mir- 197 had regulatory connections when it was lost during the diversification of the rodents.

### miRNAs are lost in tandem with TFs

Our results, thus far, provide a first assessment of the likely consequences of the multiple miRNA losses on the gene regulatory network during the *Eumuroidea* diversification. We next investigated whether this change of gene regulation on the post-transcriptional regulation was accompanied by a corresponding (and hitherto overlooked) change at the transcriptional level. To this end, the phylogenetic profiles of human miRNAs were supplemented with the phylogenetic profiles of all human protein-coding genes. We identified 127 protein-coding genes, whose phylogenetic profiles indicated that they were co-lost with the 15 miRNA families (**Error! Reference source not found.**, Figure 5A). We again ensured that the alleged loss does not represent sensitivity limits of the ortholog search (Supplementary Figure S7, Supplementary Figure S8). Additionally, there is no overlap of primary targets or miRTarBase targets with the lost protein-coding genes, yielding no evidence that the miRNA loss followed a loss of their respective target genes (Supplementary Figure S7).

**Figure 5:**
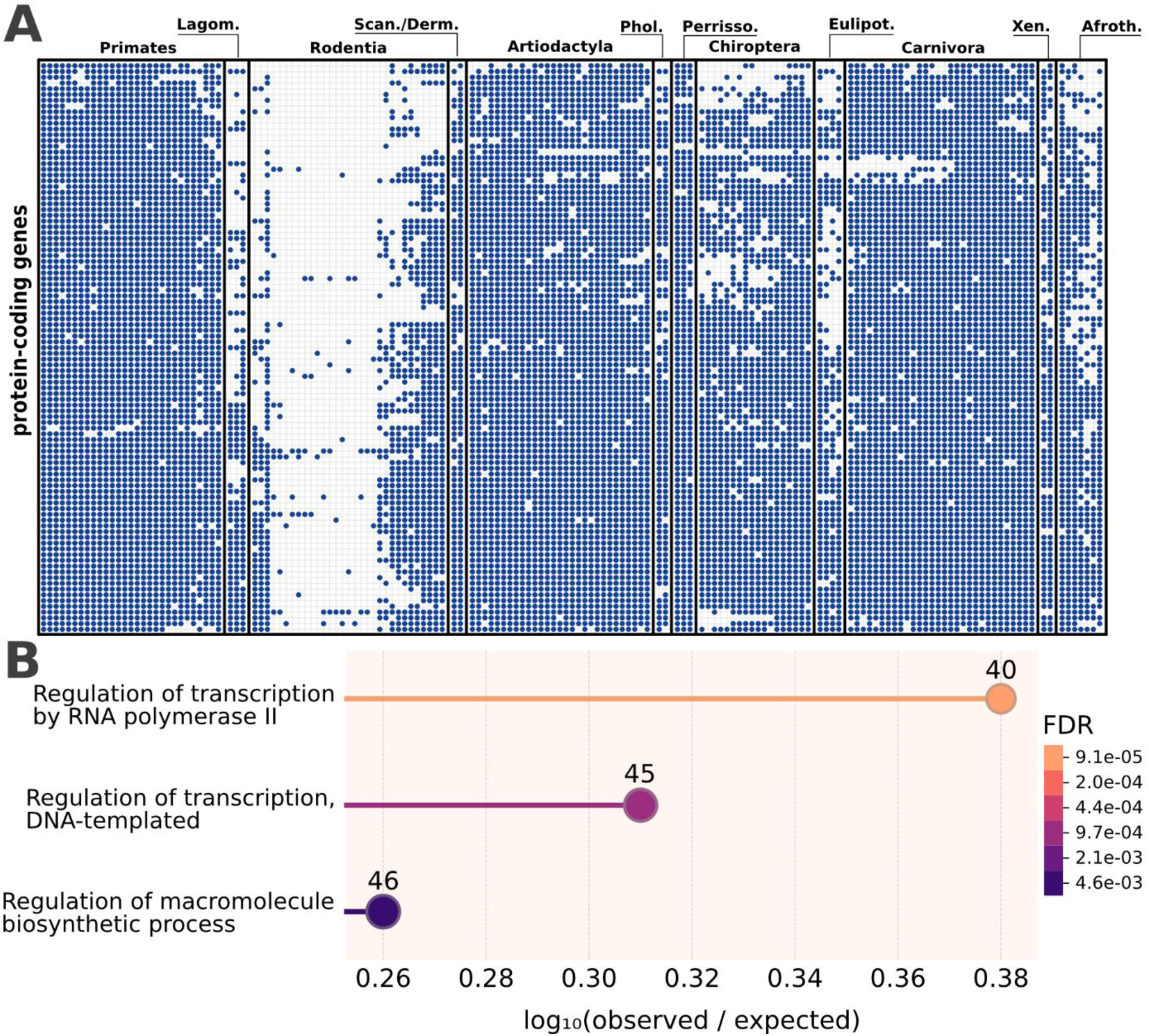
Protein-coding genes lost in tandem with the miRNAs are enriched for transcription-related biological processes. A) Phylogenetic profiles of orthologs to human protein-coding genes. The presence of an ortholog is indicated by a dot. B) Enrichment analysis for the “Biological Process” class, performed by the STRING database (**37**). The number of genes annotated with the respective GO-term are shown above the node.

A GO-term enrichment analysis revealed a significant overrepresentation of genes involved in the regulation of transcription (Figure 5B). In line with this, the set contained 37 TFs listed in the Human Transcription Factor database (HTFD) (36), which is significantly more than expected by chance (Fisher’s exact test, 4.2-fold enrichment, p = 3.8E-15). We next collected available TF-gene interactions for the 37 lost TFs to shed further light on the likely functional consequences of these gene losses. Five TFs were represented in the RegNetwork database of TF-gene interactions (38) and three were represented in the text-mining database of regulatory relationships TRRUST (39). For these 8 TFs, a total of 28 interactions are recorded (**Error! Reference source not found.**). Known TF target genes are enriched for the regulation of DNA repair and the regulation of transcription (Supplementary Figure S9), hinting at an even more deeply reticulated impact on the regulatory network.

To put the loss of the 37 TFs in the *Eumuroidea* into a larger context of TF evolution in mammals, we filtered the phylogenetic profiles of human protein-coding genes for entries in the HTFD. Out of the 1604 TFs, 1019 are almost ubiquitously present (> 95% of investigated taxa). The remaining TFs have been lost on individual evolutionary lineages (**Error! Reference source not found.**), and the m ost pronounced loss happened in the *Eumuroidea* (Supplementary Figure S10). We therefore conclude that the *Eumuroidea* are unique in the extent with which their regulatory network was restructured on both TF- and miRNA-level compared to other mammalian lineages.

## Discussion

After their stable integration into a regulatory network, miRNA genes are rarely lost. Here, we have shown that 15 miRNA families are missing from contemporary *Eumuroidea* due to multiple, independent gene losses. We found that this coincides with a pronounced loss of TFs on the same evolutionary lineage. This points to a shift in gene regulation at both transcriptional and post-transcriptional levels in the taxonomic group that includes the model organisms rat and mouse.

### Functional integration of miRNAs

While a concerted loss suggests functional integration for protein-coding genes (11), this has yet to be demonstrated for miRNAs. Next to the functional integration hypothesis, it is conceivable that the 15 miRNA families were lost in the *Eumuroidea* because they had only a limited impact on the regulatory network. miRNAs that are specific to mammals were previously found to only have 11 conserved targets on average (1). Additionally, their target sites appear to be less conserved across species compared to those of older miRNAs, suggesting that the corresponding regulatory network is still considerably plastic (40). However, the rarity with which a gene is lost typically reflects its biological importance (41). A model where novel miRNAs exert little biological function (5) implies a frequent loss of miRNA genes that is not observed (16,18). Additionally, previously analyses have not considered cooperative effects of multiple younger miRNAs. Consequently, the functional relevance of mammalian miRNAs on the regulatory network might have been underestimated in previous studies. However, among mammals Eumuroidea stand out by having an accelerated evolutionary rate (42,43). This increases the probability that miRNAs with only a few regulatory connections are lost. Consequently, the accumulation of miRNA losses in *Eumuroidea* could be due to chance instead of indicating their functional integration. However, the adjusted evolutionary distances between eumuroidean species and human are not larger than, for example, the ones between human and afrotherians (44) where no pronounced miRNA losses are observed (Figure 1; 16,18). This makes it unlikely that the accumulation of miRNA losses in the Eumuroidea is a result of their fast evolutionary rate.

miRNA losses are rare, suggesting their general functional relevance. Upon the knockout of individual genes, the phenotypic effects are more pronounced for older miRNAs than for younger ones (8). This implies that older miRNAs play a more fundamental role when considering single genes. However, potential cooperative effects of younger miRNAs that emerge only when knocking out several miRNAs have not yet been extensively studied. Here, we find an initial indication of cooperative effects of co-absent miRNA families via functional overlap of their targets. However, experimentally validated targets were available for only eight out of fifteen miRNA families, limiting the comprehensiveness of the functional enrichment analysis. Despite this, we find a significant enrichment of GO terms related to developmental processes. Although the overrepresented terms remain general, they align with the finding that the expression patterns of half of human development-associated genes differ from those of their murine orthologs (45).

### Reconstructing ancestral regulatory connections

Assessing the functional impact of the miRNA co-absence in the *Eumuoridea*, necessitates the reconstruction of the ancestral state of the gene regulatory network before the rodent diversification. For this reconstruction, we identified targets of Mir-197 and Mir-769 in human and murine iPSCs. A common approach to identify miRNA targets is integrating measured changes in transcript numbers with *in silico* target prediction (46). This improves accuracy, as target prediction alone infers many false positives and typically does not account for tissue-specific dosage effects of miRNA and mRNA transcript abundance (47). Notably, some methods for predicting miRNA targets are based on the evolutionary conservation of target sites (1). Alternative approaches like the context++-score implemented in TargetScan 7 take conservation-agnostic features of the miRNA and transcript sequence into account (2). While the former approach is most likely not suitable for detecting target sites of non-native miRNAs, we show that the latter is able to identify physiologically relevant target sites of human miRNAs in murine iPSCs.

To minimize false positive target predictions, we apply strict thresholds on both minimum expression change and target prediction score. We are therefore likely to underestimate the true number of ancestral miRNA targets, and future studies may reconstruct a more comprehensive set of targets, e.g. by integrating RNA-seq with PRO-seq data (48). Notably, miRNA-transcript interactions evolve rapidly (40). Hence, the ancient regulatory connections we report have likely remained detectable after the miRNA was lost because their target sites were preserved by additional factors, such as secondary structure constraints (49). This aligns with our observation that some target sites in the ancestral targets are older than the miRNA itself (see Fig. 4B). This requirement likely limits the number of ancestral regulatory connections that can be reconstructed, even with more sophisticated methods like PRO-seq.

### Restructuring of the eumuroidean regulatory network

The *Eumuroidea* lost 15 miRNA families together with 37 TFs, which are otherwise widely conserved in mammals. Of all investigated mammalian lineages, this is by far the most striking loss of regulators of gene expression, both on the transcriptional- and post-transcriptional level (Supplementary Figure S10, (16). On first sight, this multi-level modification suggests an additive if not multiplicative effect on gene expression. However, the opposite might be true. The loss of regulators on transcriptional as well as post-transcriptional level could be compensatory. In this scenario, the miRNA families are lost to balance the effect of a prior TF loss, or vice versa. According to this ‘compensation hypothesis’, the global transcriptome of eumuroidean species should be comparable to that of other mammals. Available data does not seem to support this assumption. A study of mammalian gene expression found the global transcriptome of rabbit, which branched of prior to the diversification of the rodents, to be more similar to human than that of rat and mouse (50). Similarly, the transcriptomes of human and canine natural killer cells are more alike than those of human and mouse (51). Both studies are therefore indicating that the losses of miRNAs and of TFs observed by us correlate with a considerable remodelling of the gene regulatory network in the *Eumuroidea*. Collectively, these findings support the general notion that mice represent a suboptimal host to model human diseases. In addition to the different size, life expectancy, metabolic activity, nutrition and microbiome of the two organisms (52), they also imply altered gene regulation principles. With this study, we have begun to investigate potential drivers of gene expression changes in the *Eumuroidea*. Given that only 8 out of the 37 TFs have recorded interactions in public databases, more studies are still needed for a comprehensive functional characterization of the lost TFs.

## Conclusions

miRNAs are seldomly lost (15,16,18), directing attention to the absence of 15 miRNA families in the *Eumuroidea* lineage. Remnants of the ancient regulatory network shaped by the lost miRNAs indicate a regulatory shift that is specific to this lineage. This shift of the regulatory network extends to the regulation of gene transcription where at least 37 transcription factors were lost. We show that the *Eumuroidea* are unique in the extent with which their regulatory network was restructured compared to other mammals. Additionally, we find no evidence that this shift was functionally compensated. These findings have important implications for studies that use mouse or rat as model organisms to analyze gene expression, for example in the context of investigating human diseases. Depending on the research question, it should be considered whether a non-eumuroidean model organism like rabbit, dog or cattle is more appropriate for the comparison with human than mouse or rat.

## Data availability

The RNA-Seq datasets have been deposited and are available at NCBI GEO. The accession number for human is GSE281793 (https://www.ncbi.nlm.nih.gov/geo/query/acc.cgi?acc=GSE281793) and the mouse dataset is available with the accession number GSE281795 (https://www.ncbi.nlm.nih.gov/geo/query/acc.cgi?acc=GSE281795). Data analysis scripts are available at https://github.com/felixlangschied/rodent_loss.

## Author contributions

Felix Langschied (Conceptualization [equal], Data Curation [equal], Formal Analysis [lead], Investigation [equal], Software [lead], Validation [equal], Visualization [lead], Writing – Original Draft Preparation [lead], Writing – Review & Editing [equal]), Matthias Leisegang (Conceptualization [supporting], Data Curation [equal], Formal Analysis [supporting], Investigation [equal], Validation [equal], Visualization [supporting], Writing – Review & Editing [supporting]), Stefan Günther (Data Curation [equal], Formal Analysis [supporting], Investigation [supporting]), Fabian Hahne (Investigation [supporting]), Ralf P. Brandes (Conceptualization [equal], Funding Acquisition [equal], Resources [equal], Supervision [supporting], Validation [equal], Writing – Review & Editing [supporting]), Ingo Ebersberger (Conceptualization [equal], Funding Acquisition [equal], Resources [equal], Supervision [lead], Validation [equal], Writing – Original Draft Preparation [supporting], Writing – Review & Editing [equal])

## Acknowledgments

We thank Judit Izquierdo Ponce and Manuela Späth for excellent technical assistance.

## Supplementary Figures

**Supplementary Figure S1:**
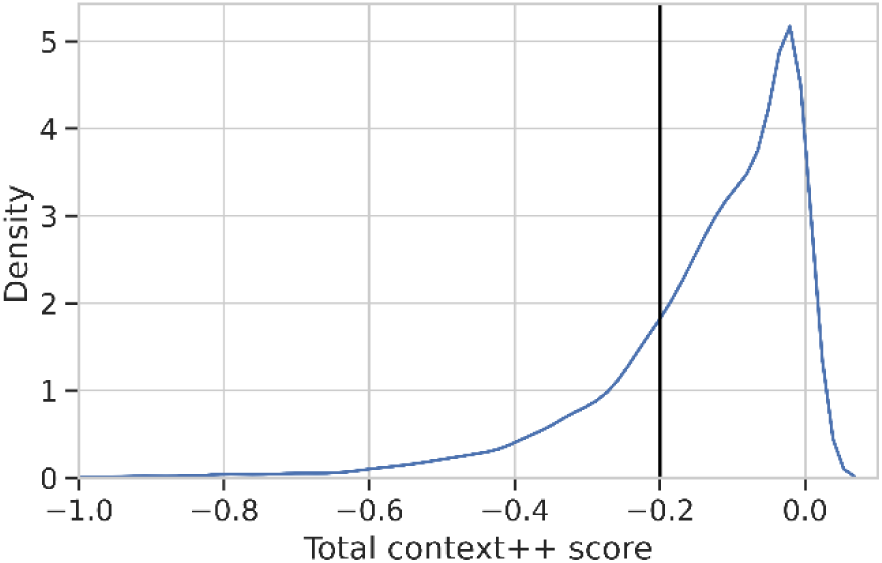
Distribution of total context++ scores of miRNA targets predicted with TargetScan7. The minimum score threshold used for finding primary miRNA targets of -0.2 is marked with a black line.

**Supplementary Figure S2:**
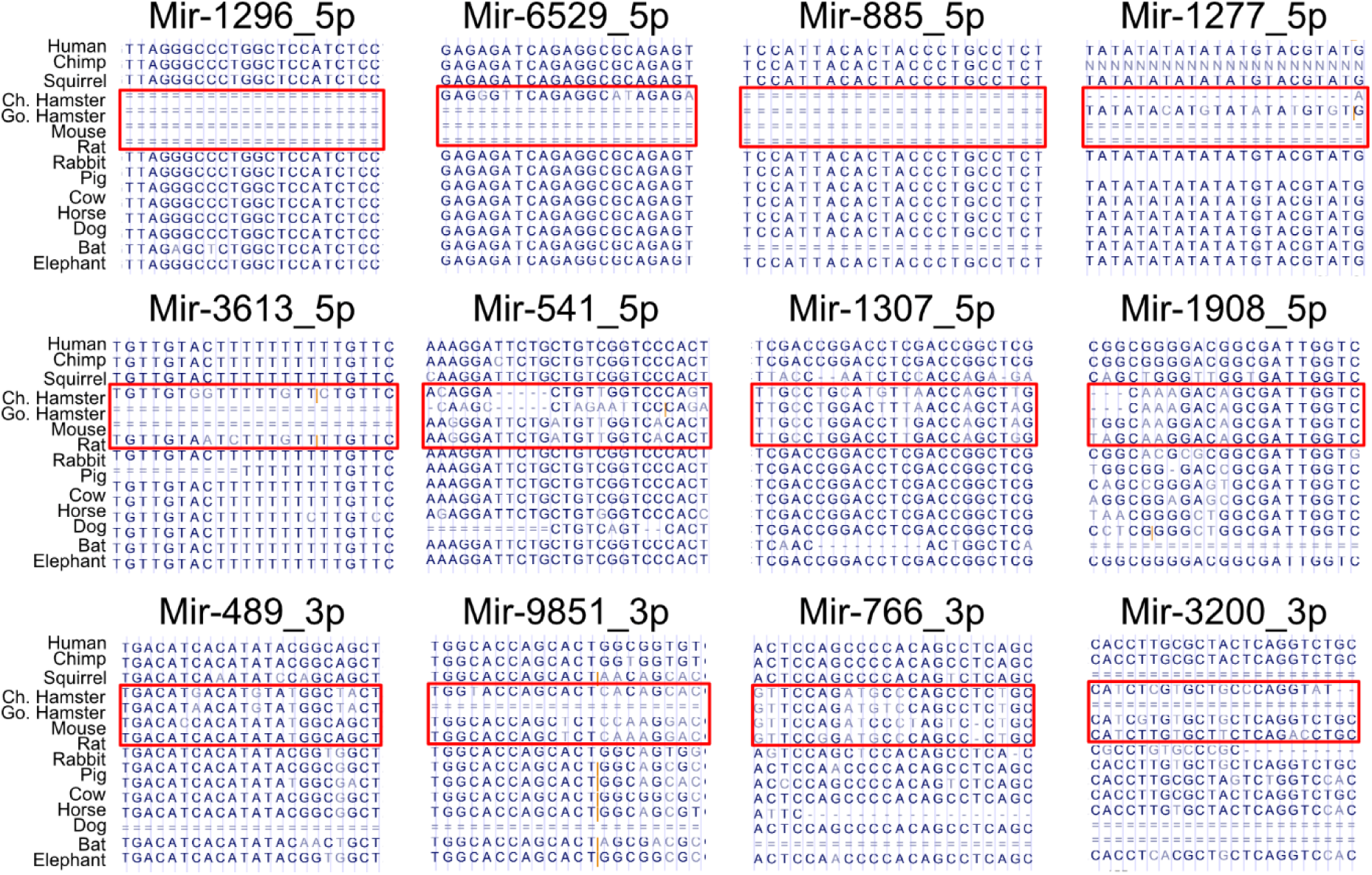
14-species whole-genome alignments of miRNA families lost the Eumuroidea. Eumuroidean species are marked with a red box.

**Supplementary Figure S3:**
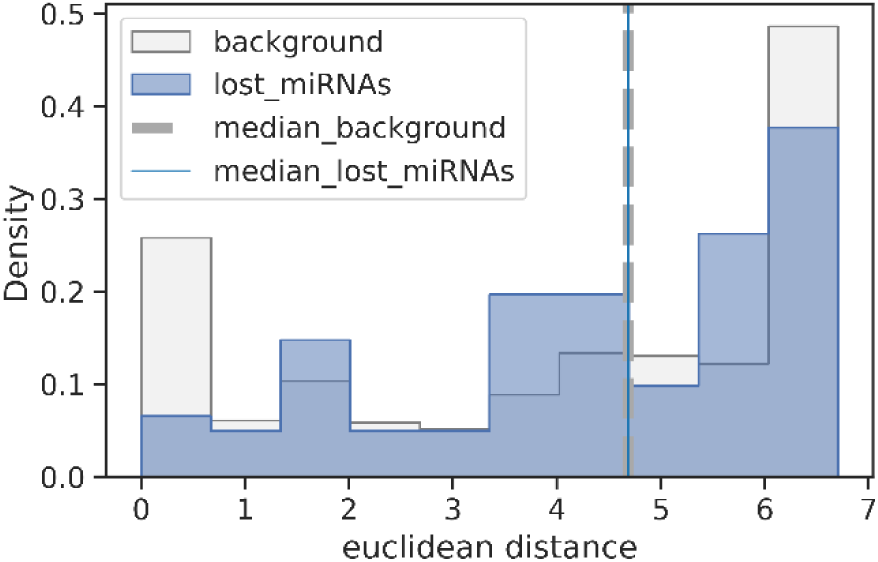
Pairwise distances of miRNA expression vectors across 45 human tissues. Normalized reads of miRNA expression were downloaded from MirGeneDB 2.1 (34). A minimum threshold of 15 was set to determine presence or absence of a miRNA in a tissue. If a miRNA family was represented by multiple genes, the gene with the highest expression in the family was chosen as representative.

**Supplementary Figure S4:**
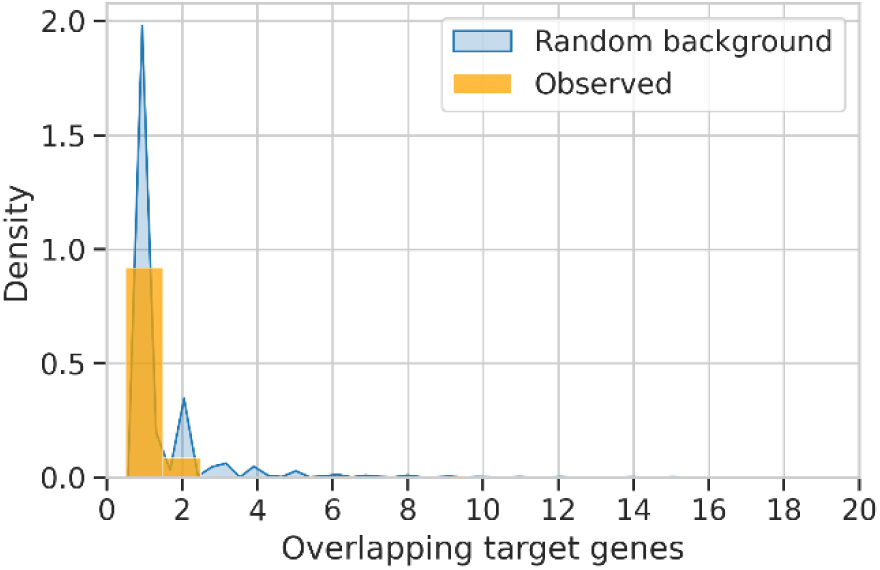
Overlap of miRTarBase targets of lost miRNAs.

**Supplementary Figure S5:**
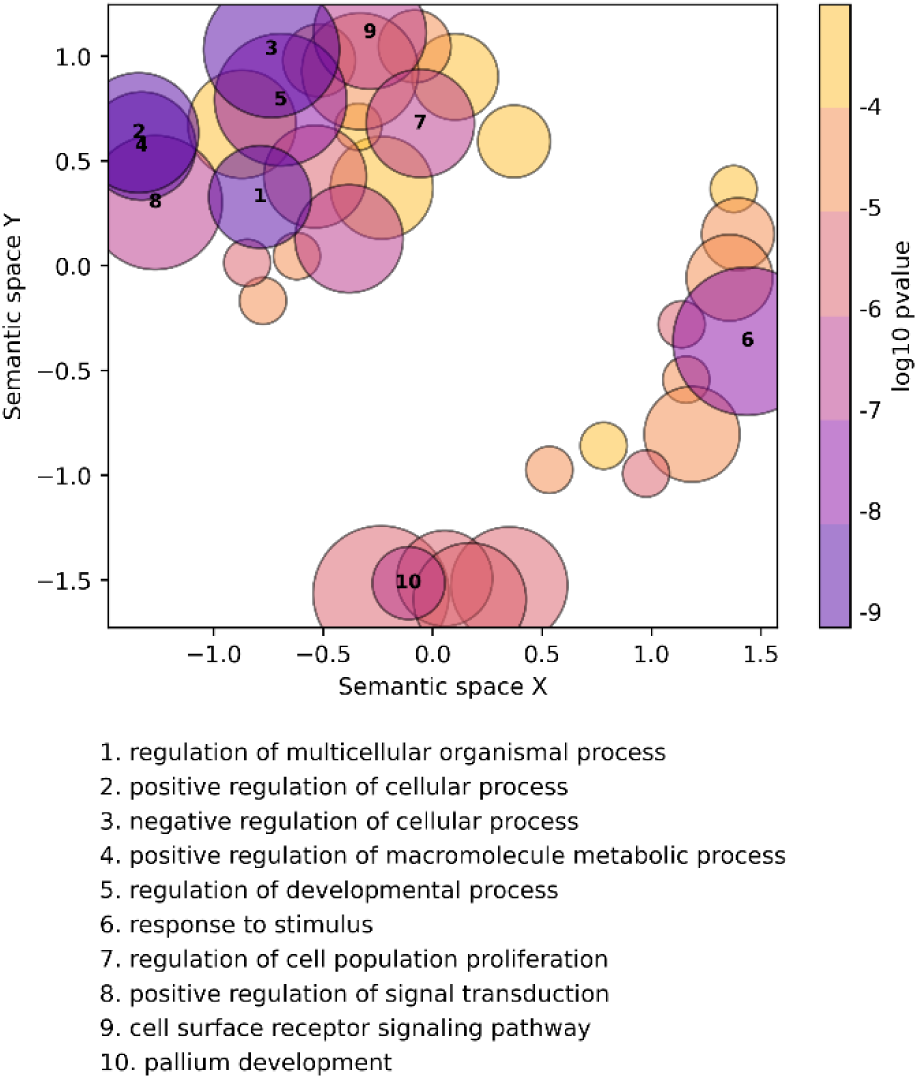
Significantly enriched GO-terms of miRTarBase targets of lost miRNAs. Visualized using semantic clustering performed by GO-Figure (53).

**Supplementary Figure S6:**
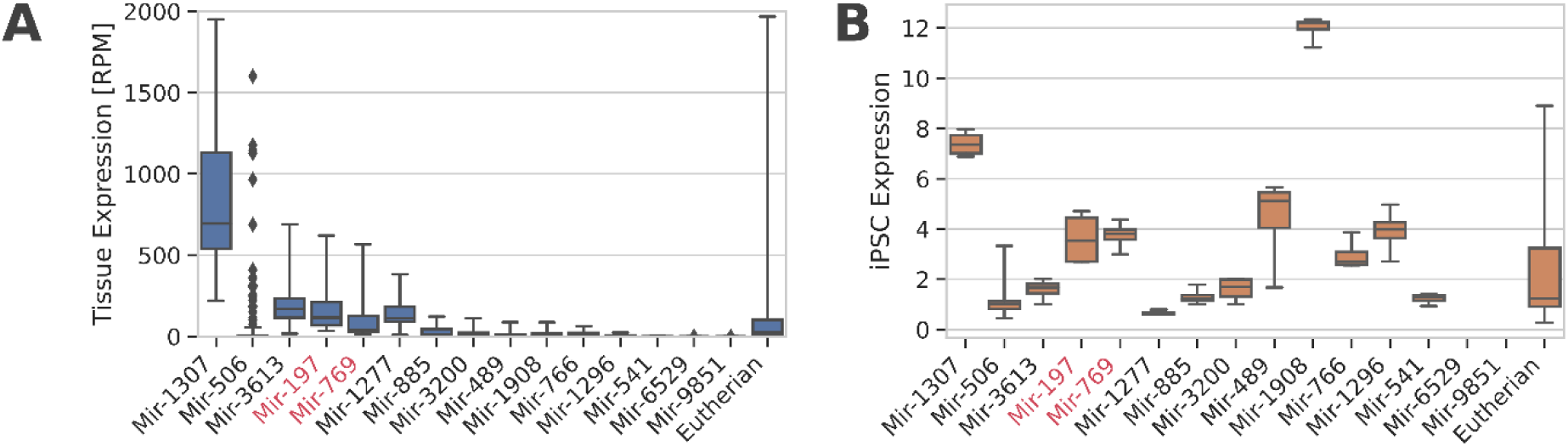
Expression of miRNA genes in human tissues and human iPSCs from the 15 families of interest and of all miRNAs gained in the LCA of Eutherians. A) RNAseq data of 45 tissues from MirGeneDB (34). B) Microarray expression levels in human iPSCs reported by (54).

**Supplementary Figure S7:**
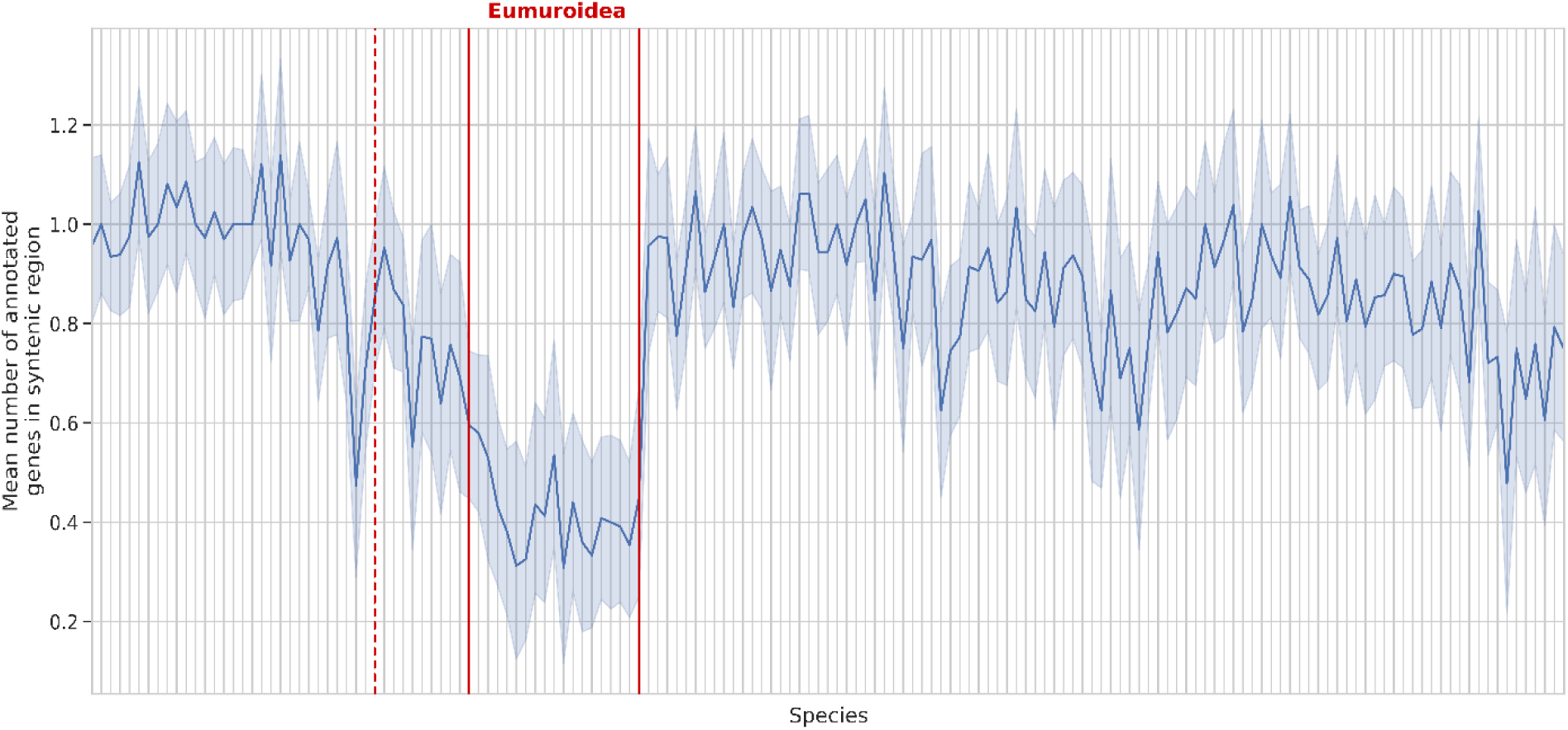
Number of annotated genes in genomic region that is shared syntenic to human reference proteins. First, the number of annotated genes in all identified syntenic regions was counted. In taxa where no ortholog was found (i.e. the Eumuroidea), this value drops below 1, indicating that there is no annotated gene in the respective region.

**Supplementary Figure S8:**
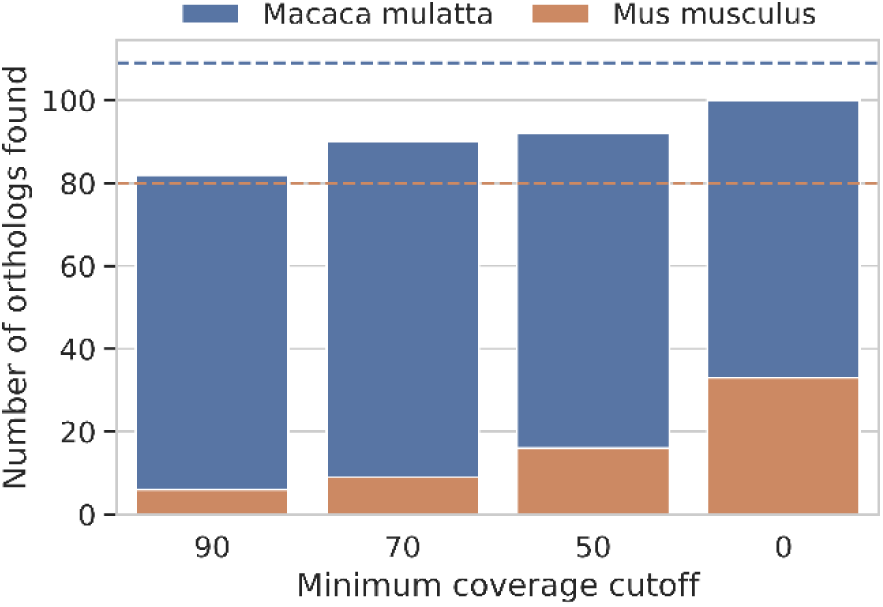
Sequence similarity search identifies ruins of lost genes. A tBLASTn search of the human protein-sequence was performed in the respective shared syntenic region of the mouse genome. Number of identified syntenic region per species are marked with a dashed line. This identified short fragments that share a significant sequence similarity to the lost genes. Thus, the syntenic regions in mouse still harbour remnants of the lost genes but they are sufficiently degraded so that no gene was inferred in the region. This strongly indicates that the genes were indeed lost (55), and that the functions encoded by them are most likely absent in the Eumuroidea.

**Supplementary Figure S9:**
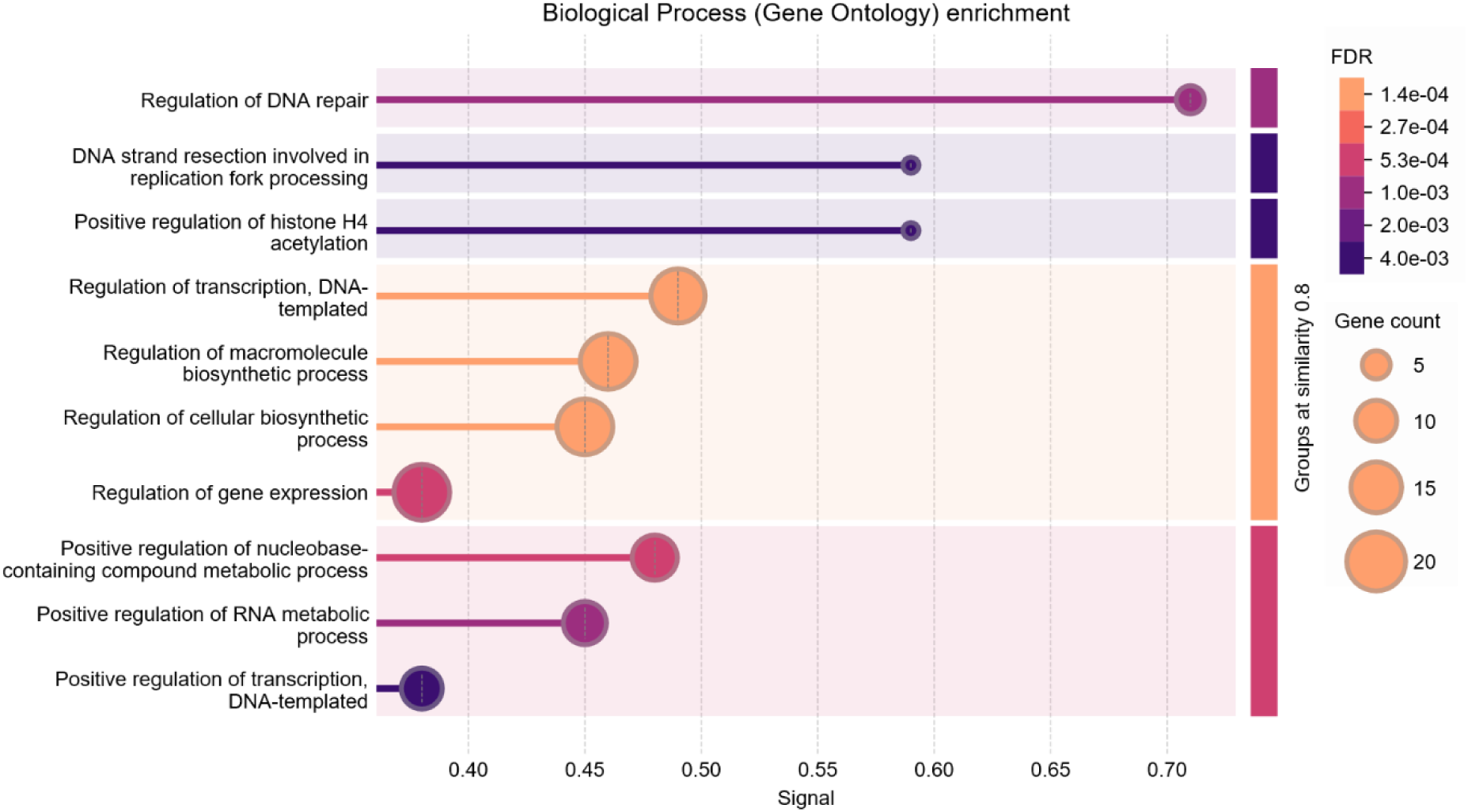
GO-Enrichment analysis of the genes targeted by TFs lost in the Eumuroidea. Performed by the STRING database (37).

**Supplementary Figure S10:**
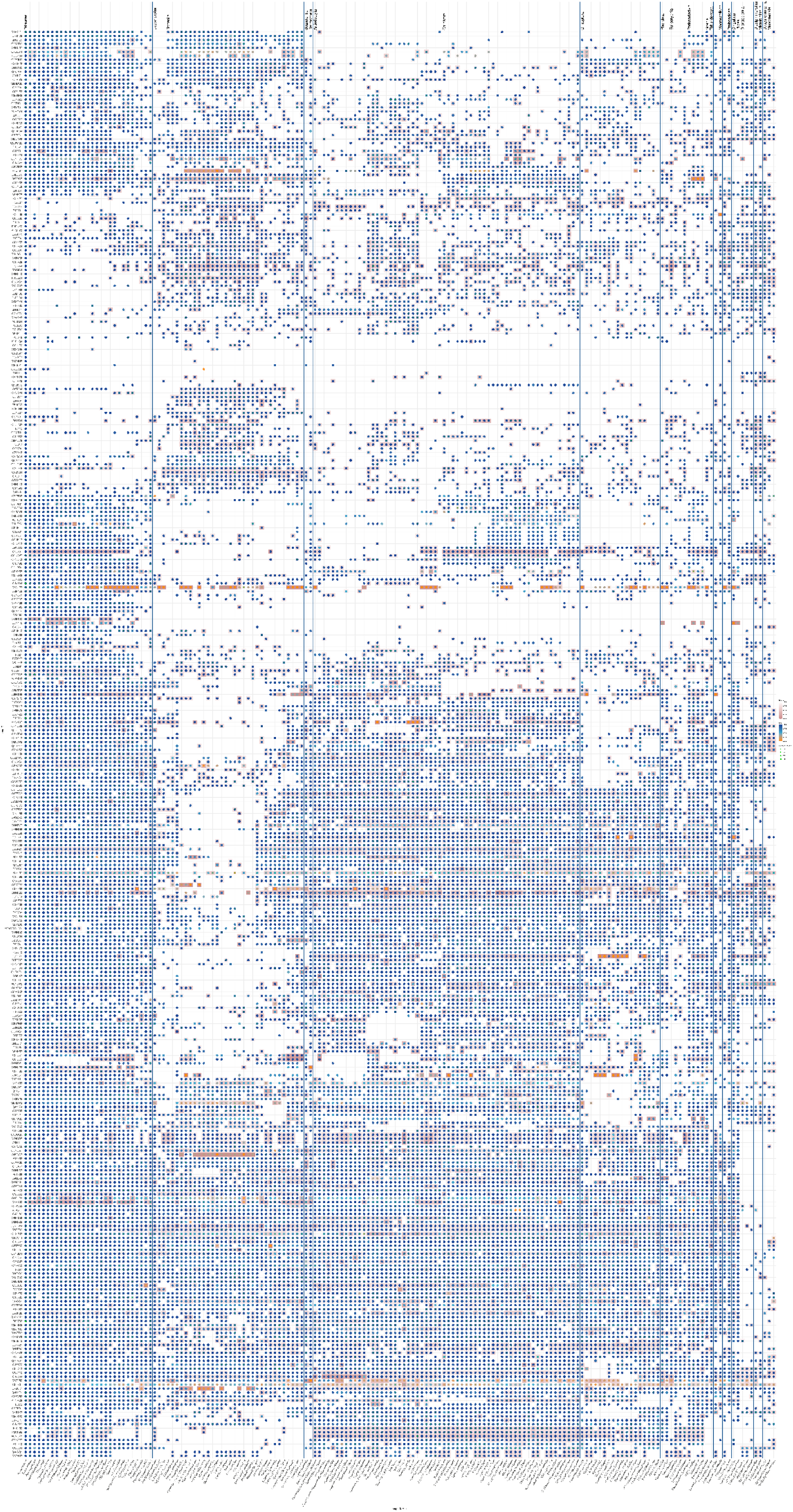
Phylogenetic Profile of non-conserved human transcription factors.

## References

1. Friedman RC, Farh KKH, Burge CB, Bartel DP. Most mammalian mRNAs are conserved targets of microRNAs. Genome Res. 2009 Jan;19(1):92–105.

2. Agarwal V, Bell GW, Nam JW, Bartel DP. Predicting effective microRNA target sites in mammalian mRNAs. Izaurralde E, editor. eLife. 2015;4:e05005.

3. Lewis BP, Shih I hung, Jones-Rhoades MW, Bartel DP, Burge CB. Prediction of Mammalian MicroRNA Targets. Cell. 2003 Dec 26;115(7):787–98.

4. Nozawa M, Fujimi M, Iwamoto C, Onizuka K, Fukuda N, Ikeo K, et al. Evolutionary Transitions of MicroRNA-Target Pairs. Genome Biol Evol. 2016 Jun 4;8(5):1621–33.

5. Chen K, Rajewsky N. The evolution of gene regulation by transcription factors and microRNAs. Nature Reviews Genetics. 2007 Feb 1;8(2):93–103.

6. Iwama H, Kato K, Imachi H, Murao K, Masaki T. Human microRNAs preferentially target genes with intermediate levels of expression and its formation by mammalian evolution. PLoS One. 2018;13(5):e0198142.

7. Patel VD, Capra JA. Ancient human miRNAs are more likely to have broad functions and disease associations than young miRNAs. BMC Genomics. 2017 Aug 31;18(1):672.

8. Bartel DP. Metazoan MicroRNAs. Cell. 2018 Mar 22;173(1):20–51.

9. Yang K, Yu B, Cheng C, Cheng T, Yuan B, Li K, et al. Mir505-3p regulates axonal development via inhibiting the autophagy pathway by targeting Atg12. Autophagy. 2017 Oct 3;13(10):1679–96.

10. Marty V, Labialle S, Bortolin-Cavaillé ML, Ferreira De Medeiros G, Moisan MP, Florian C, et al. Deletion of the miR-379/miR-410 gene cluster at the imprinted Dlk1-Dio3 locus enhances anxiety-related behaviour. Hum Mol Genet. 2016 Feb 15;25(4):728–39.

11. Pellegrini M, Marcotte EM, Thompson MJ, Eisenberg D, Yeates TO. Assigning protein functions by comparative genome analysis: protein phylogenetic profiles. Proc Natl Acad Sci U S A. 1999 Apr 13;96(8):4285–8.

12. Dembech E, Malatesta M, De Rito C, Mori G, Cavazzini D, Secchi A, et al. Identification of hidden associations among eukaryotic genes through statistical analysis of coevolutionary transitions. Proc Natl Acad Sci U S A. 2023 Apr 18;120(16):e2218329120.

13. van Hooff JJ, Tromer E, van Wijk LM, Snel B, Kops GJ. Evolutionary dynamics of the kinetochore network in eukaryotes as revealed by comparative genomics. EMBO Rep. 2017 Sep;18(9):1559– 71.

14. Djahanschiri B, Di Venanzio G, Distel JS, Breisch J, Dieckmann MA, Goesmann A, et al. Evolutionarily stable gene clusters shed light on the common grounds of pathogenicity in the Acinetobacter calcoaceticus-baumannii complex. PLoS Genet. 2022 Jun;18(6):e1010020.

15. Fromm B, Billipp T, Peck LE, Johansen M, Tarver JE, King BL, et al. A Uniform System for the Annotation of Vertebrate microRNA Genes and the Evolution of the Human microRNAome. Vol. 49, Annual Review of Genetics. Annual Reviews; 2015. p. 213–42.

16. Langschied F, Leisegang MS, Brandes RP, Ebersberger I. ncOrtho: efficient and reliable identification of miRNA orthologs. Nucleic Acids Res. 2023 Jul 21;51(13):e71.

17. Fromm B, Worren MM, Hahn C, Hovig E, Bachmann L. Substantial Loss of Conserved and Gain of Novel MicroRNA Families in Flatworms. Molecular Biology and Evolution. 2013 Dec 1;30(12):2619– 28.

18. Tarver JE, Taylor RS, Puttick MN, Lloyd GT, Pett W, Fromm B, et al. Well-Annotated microRNAomes Do Not Evidence Pervasive miRNA Loss. Genome Biol Evol. 2018 Jun 1;10(6):1457–70.

19. Nawrocki EP, Eddy SR. Infernal 1.1: 100-fold faster RNA homology searches. Bioinformatics. 2013 Nov 15;29(22):2933–5.

20. Fromm B, Domanska D, Høye E, Ovchinnikov V, Kang W, Aparicio-Puerta E, et al. MirGeneDB 2.0: the metazoan microRNA complement. Nucleic Acids Res. 2020 Jan 8;48(D1):D132–41.

21. O’Leary NA, Wright MW, Brister JR, Ciufo S, Haddad D, McVeigh R, et al. Reference sequence (RefSeq) database at NCBI: current status, taxonomic expansion, and functional annotation. Nucleic Acids Res. 2016 Jan 4;44(D1):D733–745.

22. Tran V, Langschied F, Muelbaier H, Dosch J, Arthen F, Balint M, et al. Tracing the taxonomic distribution of plant cell wall degrading enzymes across the tree of life using feature architecture aware orthology assignments. bioRxiv. 2024 Jan 1;2024.10.16.618745.

23. Huang HY, Lin YCD, Cui S, Huang Y, Tang Y, Xu J, et al. miRTarBase update 2022: an informative resource for experimentally validated miRNA–target interactions. Nucleic Acids Research. 2022 Jan 7;50(D1):D222–30.

24. Csűös M. Count: evolutionary analysis of phylogenetic profiles with parsimony and likelihood. Bioinformatics. 2010 Aug 1;26(15):1910–2.

25. Hahner F, Moll F, Warwick T, Hebchen DM, Buchmann GK, Epah J, et al. Nox4 promotes endothelial differentiation through chromatin remodeling. Redox Biol. 2022 Sep;55:102381.

26. Sommer CA, Stadtfeld M, Murphy GJ, Hochedlinger K, Kotton DN, Mostoslavsky G. Induced Pluripotent Stem Cell Generation Using a Single Lentiviral Stem Cell Cassette. Stem Cells. 2009 Mar 1;27(3):543–9.

27. Busk PK. A tool for design of primers for microRNA-specific quantitative RT-qPCR. BMC Bioinformatics. 2014 Jan 28;15(1):29.

28. Bolger AM, Lohse M, Usadel B. Trimmomatic: a flexible trimmer for Illumina sequence data. Bioinformatics. 2014 Aug 1;30(15):2114–20.

29. Dobin A, Davis CA, Schlesinger F, Drenkow J, Zaleski C, Jha S, et al. STAR: ultrafast universal RNA-seq aligner. Bioinformatics. 2013 Jan 1;29(1):15–21.

30. Broad Institute. Picard toolkit [Internet]. Broad Institute, GitHub repository. Broad Institute; 2019. Available from: https://broadinstitute.github.io/picard/

31. Liao Y, Smyth GK, Shi W. featureCounts: an efficient general purpose program for assigning sequence reads to genomic features. Bioinformatics. 2014 Apr 1;30(7):923–30.

32. Love MI, Huber W, Anders S. Moderated estimation of fold change and dispersion for RNA-seq data with DESeq2. Genome Biology. 2014 Dec 5;15(12):550.

33. Raney BJ, Barber GP, Benet-Pagès A, Casper J, Clawson H, Cline MS, et al. The UCSC Genome Browser database: 2024 update. Nucleic Acids Research. 2024 Jan 5;52(D1):D1082–8.

34. Fromm B, Høye E, Domanska D, Zhong X, Aparicio-Puerta E, Ovchinnikov V, et al. MirGeneDB 2.1: toward a complete sampling of all major animal phyla. Nucleic Acids Research. 2022 Jan 7;50(D1):D204–10.

35. Bellman R, Chen J, Chen L, Nomikou N, Tsui J, Hamilton G, et al. Comparison of gene expression between human and mouse iPSC-derived cardiomyocytes for stem cell therapies of cardiovascular defects via bioinformatic analysis. Translational Medicine Communications. 2023 Mar 17;8(1):9.

36. Lambert SA, Jolma A, Campitelli LF, Das PK, Yin Y, Albu M, et al. The Human Transcription Factors. Cell. 2018 Feb 8;172(4):650–65.

37. Szklarczyk D, Gable AL, Lyon D, Junge A, Wyder S, Huerta-Cepas J, et al. STRING v11: protein-protein association networks with increased coverage, supporting functional discovery in genome-wide experimental datasets. Nucleic acids research. 2019 Jan;47(D1):D607–13.

38. Liu ZP, Wu C, Miao H, Wu H. RegNetwork: an integrated database of transcriptional and post-transcriptional regulatory networks in human and mouse. Database. 2015 Jan 1;2015:bav095.

39. Han H, Cho JW, Lee S, Yun A, Kim H, Bae D, et al. TRRUST v2: an expanded reference database of human and mouse transcriptional regulatory interactions. Nucleic Acids Res. 2018 Jan 4;46(D1):D380–6.

40. Simkin A, Geissler R, McIntyre ABR, Grimson A. Evolutionary dynamics of microRNA target sites across vertebrate evolution. PLoS Genet. 2020 Feb;16(2):e1008285.

41. Krylov DM, Wolf YI, Rogozin IB, Koonin EV. Gene loss, protein sequence divergence, gene dispensability, expression level, and interactivity are correlated in eukaryotic evolution. Genome Res. 2003 Oct;13(10):2229–35.

42. Churakov G, Sadasivuni MK, Rosenbloom KR, Huchon D, Brosius J, Schmitz J. Rodent Evolution: Back to the Root. Molecular Biology and Evolution. 2010 Jun 1;27(6):1315–26.

43. Thybert D, Roller M, Navarro FCP, Fiddes I, Streeter I, Feig C, et al. Repeat associated mechanisms of genome evolution and function revealed by the Mus caroli and Mus pahari genomes. Genome Res. 2018 Apr;28(4):448–59.

44. Jain A, Perisa D, Fliedner F, von Haeseler A, Ebersberger I. The Evolutionary Traceability of a Protein. Genome Biol Evol. 2019 Feb 1;11(2):531–45.

45. Cardoso-Moreira M, Sarropoulos I, Velten B, Mort M, Cooper DN, Huber W, et al. Developmental Gene Expression Differences between Humans and Mammalian Models. Cell Rep. 2020 Oct 27;33(4):108308.

46. Hausser J, Zavolan M. Identification and consequences of miRNA–target interactions — beyond repression of gene expression. Nature Reviews Genetics. 2014 Sep 1;15(9):599–612.

47. Fridrich A, Hazan Y, Moran Y. Too Many False Targets for MicroRNAs: Challenges and Pitfalls in Prediction of miRNA Targets and Their Gene Ontology in Model and Non-model Organisms. Bioessays. 2019 Apr;41(4):e1800169.

48. Patel RK, West JD, Jiang Y, Fogarty EA, Grimson A. Robust partitioning of microRNA targets from downstream regulatory changes. Nucleic Acids Research. 2020 Sep 25;48(17):9724–46.

49. Mayr C. What Are 3’ UTRs Doing? Cold Spring Harb Perspect Biol. 2019 Oct 1;11(10).

50. Cardoso-Moreira M, Halbert J, Valloton D, Velten B, Chen C, Shao Y, et al. Gene expression across mammalian organ development. Nature. 2019 Jul;571(7766):505–9.

51. Gingrich AA, Reiter TE, Judge SJ, York D, Yanagisawa M, Razmara A, et al. Comparative Immunogenomics of Canine Natural Killer Cells as Immunotherapy Target. Front Immunol. 2021;12:670309.

52. Rydell-Törmänen K, Johnson JR. The Applicability of Mouse Models to the Study of Human Disease. Methods Mol Biol. 2019;1940:3–22.

53. Reijnders MJMF, Waterhouse RM. Summary Visualizations of Gene Ontology Terms With GO-Figure! Frontiers in Bioinformatics [Internet]. 2021;1. Available from: https://www.frontiersin.org/articles/10.3389/fbinf.2021.638255

54. Kulcenty K, Wroblewska JP, Rucinski M, Kozlowska E, Jopek K, Suchorska WM. MicroRNA Profiling During Neural Differentiation of Induced Pluripotent Stem Cells. International journal of molecular sciences. 2019 Jul;20(15).

55. Brawand D, Wahli W, Kaessmann H. Loss of egg yolk genes in mammals and the origin of lactation and placentation. PLoS Biol. 2008 Mar 18;6(3):e63.

